# Comparative proteomic analysis of vancomycin-sensitive and vancomycin-intermediate resistant *S. aureus*

**DOI:** 10.1101/2021.08.03.455009

**Authors:** Jia Xu, Xin Jun Han, Ya Li Yang, Pei Lan Peng, Yong Zhi Hou, Zhao Yu, Pei Ru Han, Jian Hu, Xiao Xue Ma

## Abstract

The extensive use of vancomycin has led to the development of MRSA (*methicillin-resistant Staphylococcus aureus*) strains with varying degrees of resistance to vancomycin. Accumulation of surplus cell wall material, decreased cross-linking of peptidoglycan, and/ or other cell wall alterations has been put forward to explain the VISA phenotype. To our knowledge, the protein profiles of hVISA and VISA strains are rarely analyzed via quantitative comparative proteomics. In this study, we subjected subcellular fractions isolated from two isogenic S. aureus strains (*vancomycin-intermediate resistant S. aureus*) to proteomic analysis, using an integrated quantitative proteomic approach assisted by bioinformatic analysis. In total,128 up-regulated proteins (mainly AhpC, Adh, ArgF et al.) were identified,21 down-regulated proteins (mainly LtaS, SdrD, MsrR, MsrB, OatA et al.) were obtained. The largest group of differentially expressed proteins is composed of enzymic proteins associated with metabolic and catalytic activity, which accounts for 50% and 51% of the total proteins, respectively.Some proteins which take an indispensable part in the regulatory networks of *S. aureus* with vancomycin treatment are related to Cell wall metabolism ( MurA ), Cell adhesion ( SdrC, SdrD, ClfA, ClfB ), Proteolysis ( Atl, LytM, SceD ) and Pressure response ( MsrA, MsrB, AhpC ) process.In conclusion, our proteomic study revealed regulatory proteins associated with vancomycin resistance in S. aureus, and some of these proteins are involved in the regulation of cell metabolism and function, providing protential targets for further development of drug resistance strategies.

---

*Staphylococcus aureus* is a ubiquitous bacterium responsible for both community-associated and hospital-acquired infections. It can cause a wide variety of diseases, such as skin and soft tissue infections, bloodstream infections, severe pneumonia, endocarditis, as well as toxin-mediated syndromes like toxic shock and food-borne gastroenteritis[1, 2]. A Methicillin-resistant *Staphylococcus aureus* (MRSA) was first identified in England in 1961, and then it rendered *S. aureus* a worldwide severe health threat [3].The glycopeptide antibiotic vancomycin which was first released in 1958, has been used successfully for the treatment of serious MRSA infections. Unfortunately, the high prevalence of nosocomial MRSA infections and the extesive use of vancomycin has led to the development of *S. aureus* strains with varying degrees of resistannce to vancomycin[4, 5].

In 1997, *S. aureus* with reduced susceptibility to vancomycin was reported from Japan [6]. This report described the first isolation of vancomycin-intermediate resistant *S. aureus* (VISA, MIC of 8µg/ml) from a surgical wound infection from a 4-month-old individual who had undergone cardiac surgery, where vancomycin failed to cure the infection. It may be an early warning that *S. aureus* strains with full resistance to vancomycin will emerge. Some MRSA strains have been reported that contain limited subpopulations with intermediate resistance to glycopeptides, while most of the population remains glycopeptide-susceptible[7, 8]. These strains are termed heteroresistant vancomycin-intermediate *S. aureus* (hVISA). The first hVISA strain (MIC of 4µg/ml) was isolated in 1996 from the sputum of a 64-year-old patient with MRSA pneumonia who failed vancomycin therapy [9]. Contrary to the VISA phenotype, highly vancomycin-resistant clinical *S. aureus* isolates (VRSA), having incorporated the *vanA* gene from *enterococci*, first emerged in hospitalized patients in 2002. Fortunately, to date only a few cases of VRSA have been observed, indicating that resistance is not spreading rapidly [10].

Several factors have been reported to be involved in the phenotypic characteristics of vancomycin resistance in hVISA and VISA, including thickened cell wall[11, 12],accumulation of surplus cell wall material [13], decreased cross-linking of peptidoglycan[14, 15], inactivation of penicillin-binding protein D (PBP4)[16], a decrease in the level of the muropeptide amidation[11, 15], and/ or other cell wall alterations, such as increased glycan chain length.[17] Contrary to the cleared phenotypic characteristics, the genetic mechanism of VISA is far being clear. Several transcriptional profiling studies were performed using isogenic VISA strains with different vancomycin resistance level[18]. Different VISA strains have been studied by many researchers, and altered expression of genes was reported. However, until now, no unifying molecular hypothesis has been put forward to explain the VISA phenotype. To our knowledge, the protein profiles of hVISA and VISA strains are rarely analyzed via quantitative comparative proteomics. As one of the hot spots in drug-resistant bacteria research, technical means are still awaits further development. Proteomics for studying bacteria, has its own advantages and characteristics over other techniques. At present, the pathogenic mechanism as well as drug resistance mechanism of bacteria, is not completely elucidated, however it is considered to related to some genes function, proteins function. The traditional way to identify certain proteins is to use Western blotting and EliSA measurement. but human response to those proteins associatting with bacterial resistance are poorly understood. In this case, the introduction of proteomics will undoubtedly bring new impetus to the study of bacterial drug resistance.

In this study, Here, we subjected subcellular fractions isolated from these strains to proteomic analysis and attempted to gain further insight into the molecular causes of VISA. Therefore, using an integrated quantitative proteomic approach assisted by bioinformatic analysis, we comprehensively investigated the proteome profile. Intensive bioinformatics analysis was then carried out to annotate those quantifiable targets, including protein annotation, functional classification, functional enrichment, functional enrichment based cluster analysis.

## Materials and Methods

### Bacterial Strains and Culture Conditions

In this study, we used two isogenic *S. aureus* strains. The original strain was a Methicillin sensitive *S. aureus* strain (MSSA), which was obtained from the clinical specimen sputum from a 28-year-old patient with a clinical history of chronic necrotizing pneumonia. The strain was named in Arabic number 11 in accordance with the isolation order of all the strains. The minimum inhibitory concentration (MIC) of vancomycin for this strain 11 was 1µg/ml. By continuous vancomycin selection for 45 days *in vitro*, we obtained the vancomycin-intermediated resistant derivative from strain 11, namely 11Y (VISA). 11Y can eventually grow in BHI agar with vancomycin concentration as high as 7µg/ml, and the vancomycin MIC of 11Y is 4µg/ml.

Both 11 and 11Y strains were grown under agitation at 37℃ overnight in BHI broth medium with vancomycin concentration of 1µg/ml and 7µg/ml respectively. The over-night cultures were subcultured in a 1:100 dilution of fresh BHI broth with shaking (200rpm) at 37℃ without vancomycin. Still growing strains were harvested during exponential growth at an OD_578_ of 1.0. By centrifugation at 5000g and 4℃ for 10 min, approximately 2g bacteria pellets were harvested. The bacteria cells were then washed twice with sterile 0.01M phosphate-buffered saline (PBS) and stored in -80 ℃for later use.

### Protein Extraction and Digestion

The bacterial cells were first sonicated on ice using a high intensity ultrasonic processor (Scientz) in lysis buffer (8 M urea, 1% Triton-100, 10 mM DTT, 2 mM EDTA and 0.1% Protease Inhibitor Cocktail). The remaining debris was removed by centrifugation at 20,000 g at 4 °C for 10 min. Protein concentration was determined with 2-D Quant kit (GE Healthcare) according to the manufacturer’s instructions.

For digestion, the protein solution was reduced with 10 mM DTT (Sigma) for 1 h at 37 °C and alkylated with 20 mM IAA(Sigma) for 45 min at room temperature in darkness. For trypsin digestion, the protein sample was diluted by adding 100 mM TEAB to urea concentration less than 2M. Finally, trypsin (Promega) was added at 1:50 trypsin-to-protein mass ratio for the first digestion overnight and 1:100 trypsin-to-protein mass ratio for a second 4 h-digestion. Approximately 100 μg protein for each sample was digested with trypsin for the following experiments.

### TMT Labeling and HPLC Fractionation

After trypsin digestion, peptide was desalted by Strata X C18 SPE column (Phenomenex) and vacuum-dried. Peptide was reconstituted in 0.5 M TEAB and processed according to the manufacturer’s protocol for 6-plex TMT ki(Thermo). Briefly, one unit of TMT reagent (defined as the amount of reagent required to label 100 μg of protein) were thawed and reconstituted in 24 μl ACN (Fisher Chemical). The peptide mixtures were then incubated for 2 h at room temperature and pooled, desalted and dried by vacuum centrifugation.

Next, the mixtures were fractionated into fractions by high pH reverse-phase HPLC using Agilent 300Extend C18 column (5 μm particles, 4.6 mm ID, 250 mm length). Briefly, peptides were first separated with a gradient of 2% to 60% acetonitrile in 10 mM ammonium bicarbonate pH 10 over 80 min into 80 fractions, Then, the peptides were combined into 18 fractions and dried by vacuum centrifuging.

#### LC-MS/MS Analysis

Peptides were dissolved in 0.1% FA (Fluka), directly loaded onto a reversed-phase pre-column (Acclaim PepMap 100, Thermo Scientific). Peptide separation was performed using a reversed-phase analytical column (Acclaim PepMap RSLC, Thermo Scientific). The gradient was comprised of an increase from 6% to 20% solvent B (0.1% FA in 98% ACN) over 26 min, 20% to 35% in 6 min and climbing to 80% in 4 min then holding at 80% for the last 4 min, all at a constant flow rate of 280 nl/min on an EASY-nLC 1000 UPLC system, The resulting peptides were analyzed by Q ExactiveTM hybrid quadrupole - Orbitrap mass spectrometer (ThermoFisher Scientific).

The peptides were subjected to NSI source followed by tandem mass spectrometry (MS/MS) in Q ExactiveTM (Thermo) coupled online to the UPLC. Intact peptides were detected in the Orbitrap at a resolution of 70,000. Peptides were selected for MS/MS using NCE setting as 33; ion fragments were detected in the Orbitrap at a resolution of 17,500. A data-dependent procedure that alternated between one MS scan followed by 20 MS/MS scans was applied for the top 20 precursor ions above a threshold ion count of 1.5E4 in the MS survey scan with 20.0s dynamic exclusion. The electrospray voltage applied was 2.0 kV. Automatic gain control (AGC) was used to prevent overfilling of the ion trap; 5E4 ions were accumulated for generation of MS/MS spectra. For MS scans, the m/z scan range was 350 to 1800. Fixed first mass was set as 100 m/z.

#### Database Search

The resulting MS/MS data were processed using Mascot search engine (v.2.3.0). Tandem mass spectra were searched against Uniprot *S. aureus* NCTC 8325 database. Trypsin/P was specified as cleavage enzyme allowing up to 2 missing cleavages. Mass error was set to 10 ppm for precursor ions and 0.02 Da for fragment ions. Carbamidomethyl on Cys, TMT-6plex (N-term) and TMT-6plex (K) were specified as fixed modification and oxidation on Met was specified as variable modifications. FDR was adjusted to < 1% and peptide ion score was set > 20.

#### Bioinformatics Analysis

##### GO Annotation

The Gene Ontology (GO) annotation proteome was derived from the UniProt-GOA database (www. http://www.ebi.ac.uk/GOA/). Then proteins were classified by Gene Ontology annotation based on three categories: biological process, cellular component and molecular function.

##### Domain Annotation

Identified proteins domain functional description were annotated by InterProScan (a sequence analysis application) based on protein sequence alignment method, and the InterPro domain database was used.

##### KEGG Pathway Annotation

In our article, Kyoto Encyclopedia of Genes and Genomes (KEGG) database was used to annotate protein pathway. Firstly, using KEGG online service tools KAAS to annotated protein’s KEGG database description. Then mapping the annotation result on the KEGG pathway database using KEGG online service tools KEGG mapper.

##### Subcellular Localization

we used wolfpsort a subcellular localization predication soft to predict subcellular localization. Wolfpsort an updated version of PSORT/PSORT II for the prediction of eukaryotic sequences.

#### GO/KEGG Pathway, Protein Domain and Complex Functional Enrichment Analysis

Gene Ontology (GO) term association and enrichment analysis were performed using Functional Annotation Tool of DAVID Bioinformatics Resources 6.7. Encyclopedia of Genes and Genomes (KEGG) database was used to identify enriched pathways by Functional Annotation Tool of DAVID against the background of Homo sapiens. InterPro database was researched using Functional Annotation Tool of DAVID against the background of Homo sapiens. Manually curated CORUM protein complex database for human was used for protein complex analysis.

A two-tailed Fisher’s exact test was employed to test the enrichment of the protein-containing IPI entries against all IPI proteins. Correction for multiple hypothesis testing was carried out using standard false discovery rate control methods. The GO with a corrected p-value < 0.05 is considered significant.

### GO/KEGG Pathway Functional Enrichment-Based Clustering of Protein Groups Based on Protein Quantification

For further hierarchical clustering based on different protein functional classification (such as: GO, Domain, Pathway, Complex). We first collated all the categories obtained after enrichment along with their P values, and then filtered for those categories which were at least enriched in one of the clusters with P value <0.05. This filtered P value matrix was transformed by the function x = −log10 (P value). Finally these x values were z-transformed for each functional category. These z scores were then clustered by one-way hierarchical clustering (Euclidean distance, average linkage clustering) in Genesis. Cluster membership were visualized by a heat map using the “heatmap.2” function from the “gplots” R-package. Protein interaction

The STRING database (version:11.0, https://string-db.org) was used to search for and analyze the interaction between the differential proteins.The target proteins were entered into the STRING database to obtain a PPI network file, which shows the physical and functional interactions between the the differential proteins. The Cytoscape software (version 3.7.1) Network Analyzer was used to visualize the network file. PPI networks with diferent colors and sizes were drawn to show the change in protein expression (up-regulated or down-regulated) and node degree.

### Results Integrated Strategy for Quantitative Protein Analysis

In this study, we obtained a vancomycin intermediate resistant derivative strain 11Y from an MSSA stain 11 by stepwise vancomycin selection in vitro. Strain 11Y, which vancomycin MIC is 4µg/ml, can eventually grow in BHI agar with vancomycin concentration as high as 7µg/ml. In our previous study, we found that 11Y had significantly thicked cell wall, and a reduced autolytic activity as well as reduced biofilm formation ability compared to strain 11 (data not shown). Based on the phenotypic characters of 11 and 11Y described above, we choose them for further experiment.

Firstly, the strains were cultured with a certain concentration of vancomycin and harvested according to experimental requirements. Then proteins from the bacteria were extracted and digested. Finally, TMT labeling and HPLC fractionation were performed followed by high-resolution LC-MS/MS analysis, and quantitative global proteome analysis. The general experimental strategy was illustrated in **Figure 1**.

**FIG 1.**
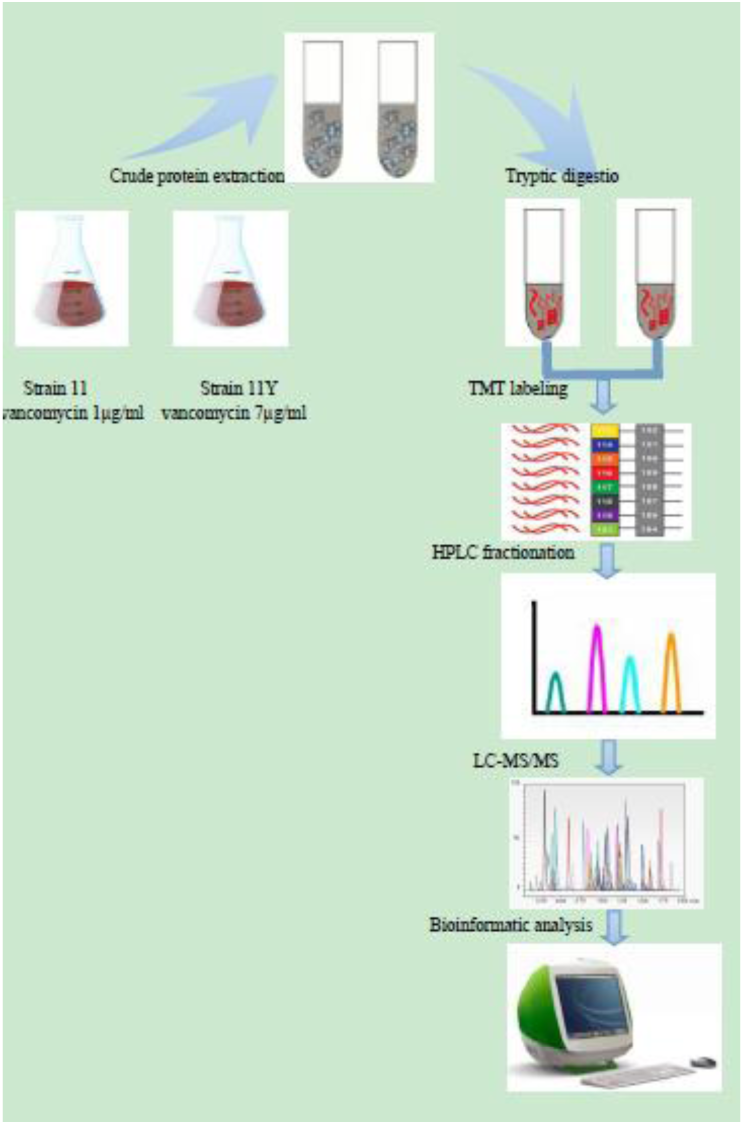
The systematic workflow of quantitative proteomics in *S. aureus*.

### Proteome QC Validation of MS Data

The MS data validation was shown in **Figure 2**. The mass error was set to 10 ppm for precursor ions and 0.02 Da for fragment ions. Firstly, we checked the mass error of all the identified peptides. The distribution of mass error is near zero and most of them are less than 0.02 Da which means the mass accuracy of the MS data fit the requirement (**Fig 2A**). After applying these parameters, we identified 1313 peptides, which exhibit different abundances depending on their length. The length of most peptides distributed between 8 and 16, which agree with the property of tryptic peptides (**Fig 2B**), that means sample preparation reach the standard.

**FIG 2.**
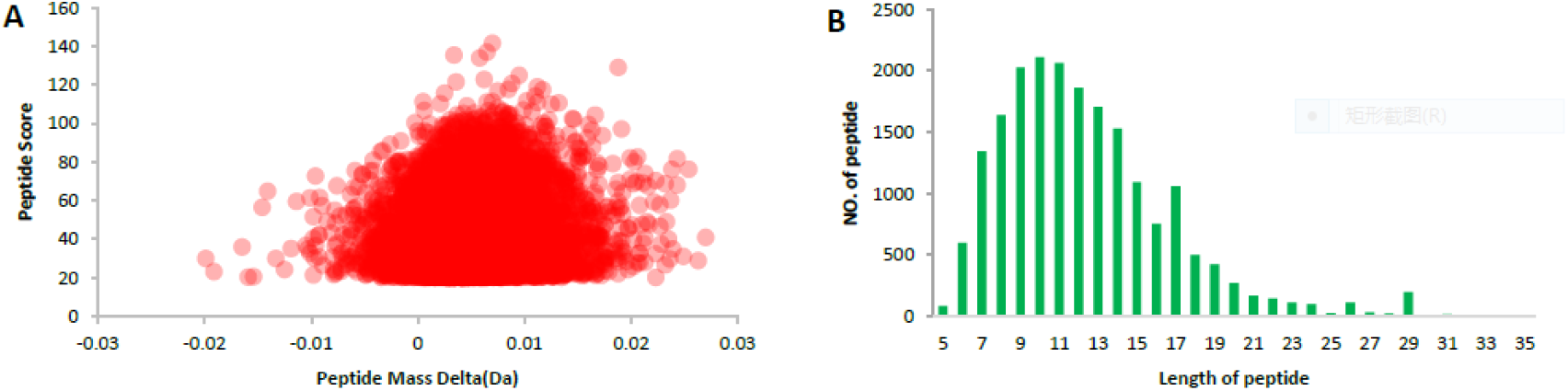
QC validation of MS data. (A) Mass error distribution of all identified peptides, (B) Peptide length distribution.

### Analysis of Differentially Expressed Protein in Strain 11 and 11Y

Further biochemistry analysis was applied to analyze the results. In total, 1313 protein groups were identified from the strains, among which 1312 proteins were quantified. When setting quantification ratio of >2.0 as up-regulated threshold, 128 up-regulated proteins (mainly AhpC, Adh, ArgF *et al*.) were identified. Setting quantification ratio of <0.5 as down-regulated threshold, 21 down-regulated proteins (mainly LtaS, SdrD, MsrR, MsrB, OatA *et al*.) were obtained. All these data were listed in appendix table A (up table), appendix table B (Down talbe). Intensive bioinformatics analysis was then carried out to annotate those quantifiable targets.

To further explore the impact of differentially expressed proteins in cell physiological process and discover internal relations between differentially expressed proteins, we classified the functions of the differentially expressed proteins and then analyzed the significance of functional enrichment including GO (biological process, cellular component, molecular function), domain, and KEGG pathway.

#### Functional Classification of Differentially Expressed Proteins

According to GO annotation information of identified protein based on their biological process, molecular function, and cellular component, we calculated the number of differentially expressed proteins in each GO term in **Table 1**. (**Fig 3**) The classification results both for biological process and molecular function showed that the largest group of differentially expressed proteins is composed of enzymic proteins associated with metabolic and catalytic activity, which accounts for 50% and 51% of the total proteins, respectively. Another large differentially expressed protein group determined by their molecular function classification comprises binding proteins, which account for 34% of all proteins. These findings indicated that the enzymic proteins associated with catalytic process and metablic process, as well as binding proteins, changed their expression levels obviously between 11 and 11Y. In terms of biological process, proteins associated with cellular processes were composed of 44% of all differentially expressed proteins. In conclude, the GO analysis suggested that proteins associated with biological process and molecular function have been most differentially expressed between strain 11 and 11Y.

**FIG 3.**
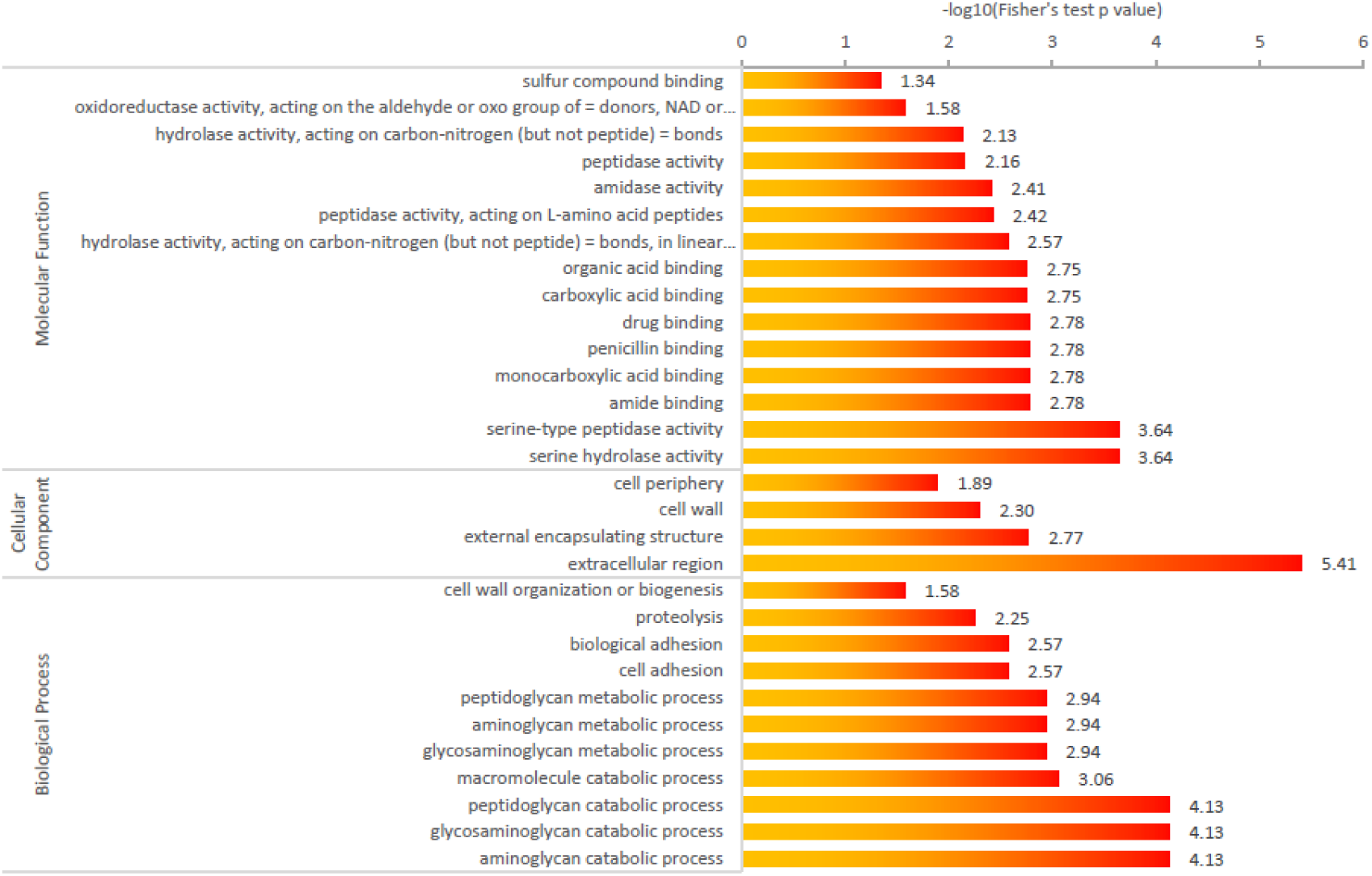
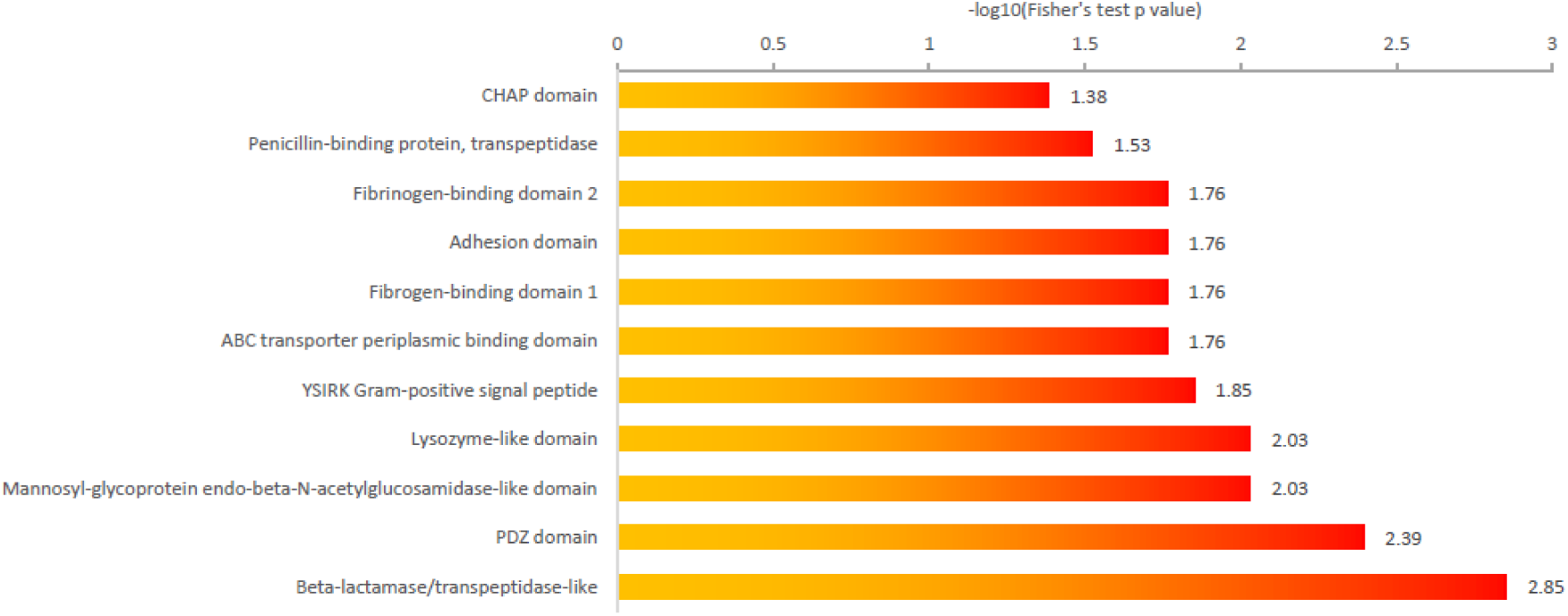
**(A)** GO-based enrichment analysis (including biological process, cellular component and molecular function). **(B)** Protein domain enrichment analysis.

**Table 1.** Differentially expressed proteins in GO terms of level 2 (11Y/11).

| GO terms level 1 | GO terms level 2 | No. of proteins | Ratio |
| --- | --- | --- | --- |
| Biological process | biological adhesion | 5 | 3.2% |
|  | localization | 5 | 3.2% |
|  | metabolic process | 50 | 32.1% |
|  | cellular component organization or | 7 | 4.5% |
|  | biogenesis | 5 | 3.2% |
|  | multi-organism process | 5 | 3.2% |
|  | response to stimulus | 2 | 1.3% |
|  | reproduction | 16 | 10.2% |
|  | single-organism process | 2 | 1.3% |
|  | signaling | 4 | 2.6% |
|  | developmental process | 44 | 28.2% |
|  | cellular process | 1 | 0.6% |
|  | cell killing | 10 | 6.4% |
|  | biological regulation |  |  |
| Molecular function | antioxidant activity | 1 | 1% |
|  | molecular transducer activity | 2 | 2% |
|  | transporter activity | 7 | 6.8% |
|  | binding | 34 | 33.3% |
|  | receptor activity | 1 | 1% |
|  | nucleic acid binding transcription factor activity | 1 | 1% |
|  | activity | 5 | 4.9% |
|  | structural molecule activity | 51 | 50% |
|  | catalytic activity |  |  |
| Cellular component | membrane | 22 | 24.4% |
|  | organelle | 6 | 6.7% |
|  | cell | 39 | 43.3% |
|  | nucleoid | 1 | 1.1% |
|  | extracellular region | 14 | 15.6% |
|  | macromolecular complex | 8 | 8.9% |

We predicted the subcellular structure and classified the differentially expressed proteins.

As shown in table 2, the subcellular location of up-regulated and down-regulated proteins were also analyzed. The results showed that most of the up-regulated proteins were distributed in the cytoplasm (16/21 proteins, 76.2%). Of the 128 down-regulated proteins, 52 proteins were located in the extracellular space (40.6%), 40 proteins were distributed in the cytoplasm (31.3%), and 36 proteins were distributed in the membrane (28.1%).

**Table 2.** Up and down-regulated proteins in subcellular location (11Y/11).

| Subcellular location | up-regulated proteins |  | down-regulated proteins |  |
| --- | --- | --- | --- | --- |
|  | No. of proteins | Ratio | No. of proteins | Ratio |
| Extracellular | 52 | 40.6% | 3 | 14.3% |
| Membrane | 36 | 28.1% | 2 | 9.5% |
| Cytoplasmic | 40 | 31.3% | 16 | 76.2% |

Gene Ontology (go) is an important bioinformatics analysis method and tool, which is used to express the attributes of genes and gene products. Go notes can be divided into three categories: biological process, cellular component and molecular function, which explain the biological function of proteins from different perspectives. We analyzed the distribution of differentially expressed proteins in go secondary annotation.

#### Functional Enrichment Analysis of Differentially Expressed Protein

Furthermore, we evaluated the enrichment function of the differentially expressed proteins in three GO categories (**Fig 3A**). Firstly, we calculate the number (a) of differentially expressed protein and the number (b) of all quantitative protein in each GO term. And then calculate the total number (c) of differentially expressed protein annotated by GO terms and the total number (d) of all quantitative protein annotated by GO terms. Finally, we use the four numbers (a, b, c, d) to calculate Fisher’ exact test P values and enrichment folds. Fold enrichment = (a/c) / (b/d).

In the biological process category, the processes related to aminoglycan catabolic process, glycosaminoglycan catabolic process, peptidoglycan catabolic process were found to be significantly enriched. This result suggested that those proteins associated with protein biosynthesis are most likely affected in *S. aureus* when 11 changed to 11Y under vancomycin stimulation. In the cellular component category, the proteins located in extracellular region were highly enriched. The result indicated that proteins composed of the extracellular region were affected a lot when the strain responsed to vancomycin treatment. In agreement with this observation, the enrichment analysis based on molecular function showed that proteins with serine-type peptidase activity and serine hydrolase activity have a higher tendency to change their expression level. When determined through the Domain Enrichment analysis shown in **Fig 3B, t**he proteins involved in Beta-lactamase/transpeptidase-like were found to be mostly enriched.

#### Enrichment-Based Clustering Analysis

For clustering analysis, the quantified proteins were divided into four quantiles (Q1-Q4). The average (x) and standard deviation (y) of the Log_10_ 11/11Y TMT ratios of all quantified peptides were calculated. The 11/11Y TMT ratio of each peptide was then transformed to a z-score based on z = (Log10Ratio – x) / y, where the Ratio is the 11/11Y TMT ratio. The cutoff Z-scores was set according to cumulative density function of normal distribution at three different percentiles 25%, 50% and 75%. Each peptide was then allocated to the quantiles based on the transformed z-score. In this way, we generated four quantiles: Q1 (0∼25%), Q2 (25∼50%), Q3 (50∼75%) and Q4 (75∼100%). Then, the enrichment -based clustering analysis (Gene Ontology, protein domain, KEGG pathway) were performed (**Fig 4A-B**).

**FIG 4.**
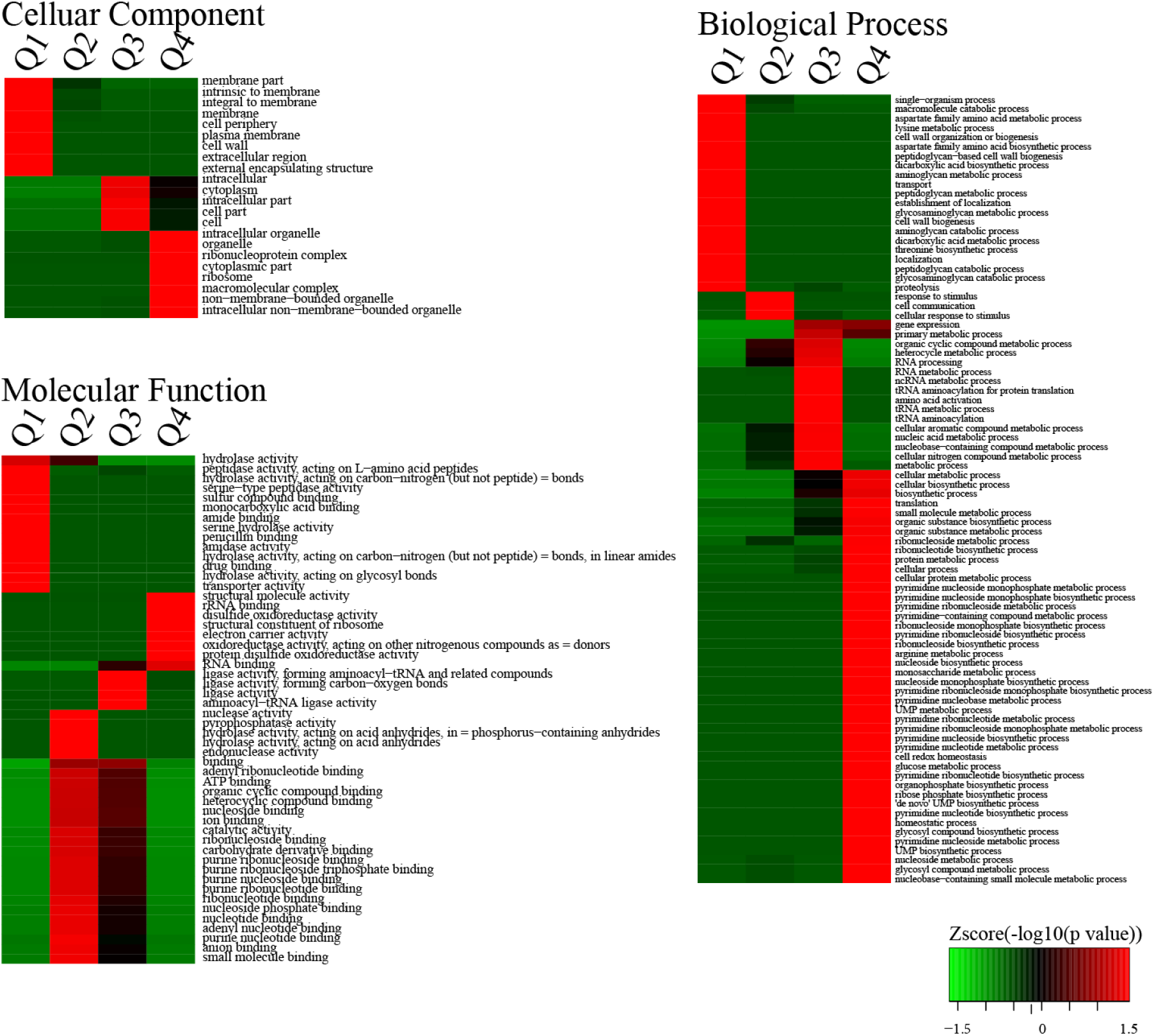
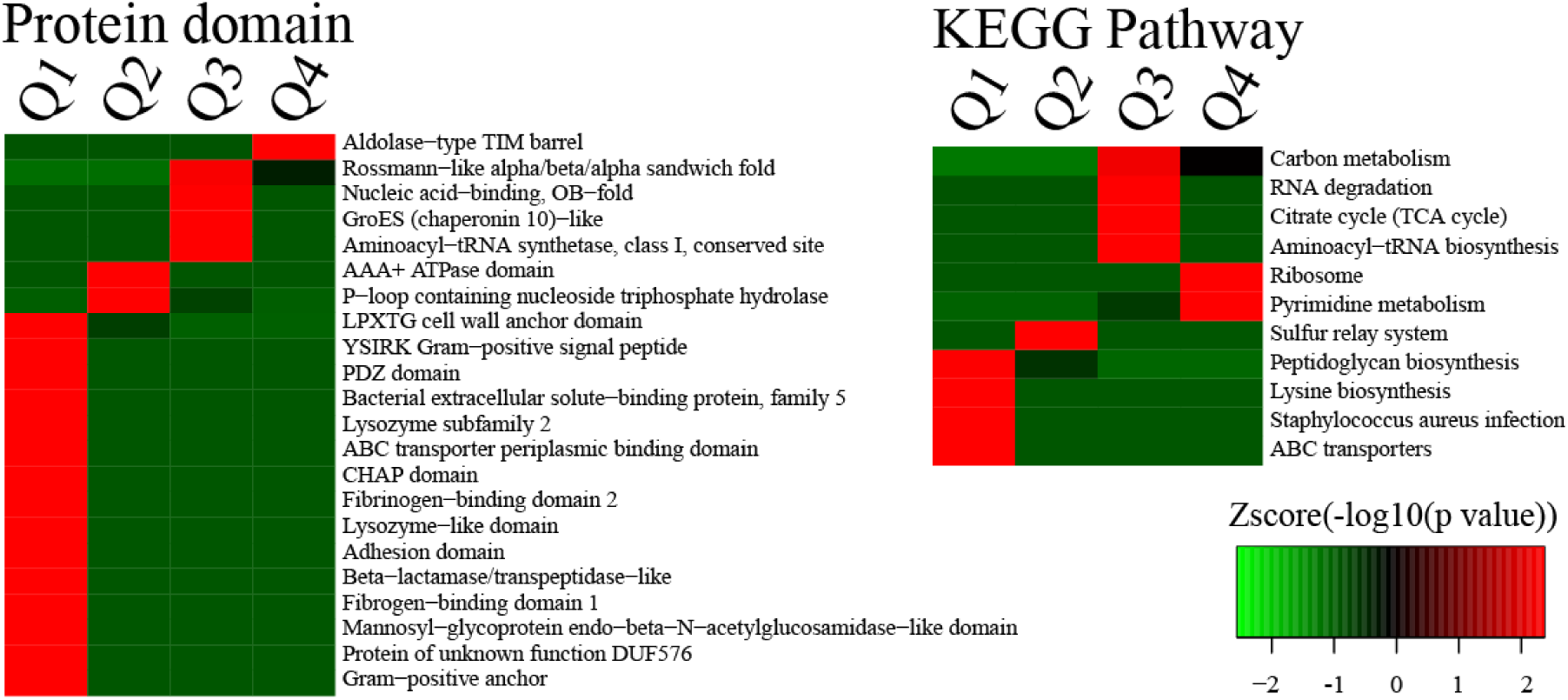
**(A)** GO enrichment-based cluster analysis. Quantifiable proteins were classified by GO annotation into three categories: cellular compartment, biological process, molecular function. In each category, the quantified proteins were divided into four quantiles according to the quantification ratio to generate four quantiles: Q1 (0∼25%), Q2 (25∼50%), Q3 (50∼75%) and Q4 (75∼100%). An enrichment analysis was performed separately in each quantile for diverse categories, and the overrepresented annotations were clustered through one-way hierarchical clustering for comparative analysis. **(B)** Protein domain and KEGG pathway enrichment-based cluster analysis.

For the cellular component analysis, we found that proteins composed of organelle, ribosome, cytoplasmic part, and macromolecular complex were highly enriched in Q4. In Q1, the proteins were focused on membrane, cell wall and external encapsulating structure. The biological process was also analyzed as showed in **Figure 4A**. A great number of proteins involved in the process of translation, organic substance biosynthesis and metabolism, nucleoside metabolism, arginine metabolic process, cell redox homeostasis, glucose metabolism were enriched in Q4. Moreover, proteins related to RNA metabolic process, tRNA metabolic process, nucleic acid metabolic process were enriched in Q3, while the proteins related to cell wall biosynthesis are rare in Q3. On the contrary, proteins focused on aspartate family amino acid metabolic process, lysine metabolic process, threonine biosynthetic process, peptidoglycan metabolic process and catabolic process, glycosaminoglycan catabolic process were enriched obviously in Q1 area.

The molecular function analysis showed that proteins with enzyme activity including hydrolase activity, peptidase activity, amidase activity, nuclease activity, endonuclease activity and so on were enriched in Q1 and Q2. In addition, binding proteins including penicillin binding, drug binding, ATP binding, ion binding were both enriched in Q1 and Q2. By contrast, the proteins enriched in Q1 and Q2 were not found in Q3 and Q4 area.

Specific domain structure is one of the major functional features in proteins. As a consequence, we next analyzed the enriched domain of those quantified proteins (**Fig 4B**). we observed that protein domains involved in ABC transporter periplasmic binding domain, fibrinogen-binding domain, lysozyme-like domain, adhesion domain, bata-lactamase/transpeptidase like domain and so on were enriched in Q1 area.

By using KEGG database, we performed a pathway enrichment-based clustering analysis of the proteome. The result showed that peptidoglycan biosynthesis pathway, lysine biosynthesis pathway, *Staphylococcus aureus* infection pathway, and ABC transport pathway were the most prominent pathways enriched in quantified proteins of vancomycin-induced derivative strain 11Y, suggesting a role of vancomycin drug pressure in these pathways. In contrast, protein expression level in the cellular pathways of carbon metabolism, RNA degradation, TCA cycle, ribosome was increased in strain 11.

## Discussion

Although specific genetic determinants of resistance mechanisms to vancomycin were identified through the use of non-proteomic approaches(e.g., van genes in vancomycin resistance)[19\20\21], recently, comparative proteomic methods provide new opportunities to understand the mechanism of antibiotic resistance. In our article, we have investigated two isogenetic *S. aureus* 11(MSSA) and 11Y(VISA) by using the novel approach to help us to analyze protein changes following th treatment of vancomycin.

In the biological process of GO enrichment, the processes related to peptidoglycan catabolism and metabolism were obviously enriched. Peptidoglycan (PG) is an essential and integral component of the bacterial cell wall found on the outside of the cytoplasmic membrane of almost all bacteria.[22–24] It is responsible for maintaining a defined cell shape, protecting the cells from osmotic lysis and serving as a scaffold for anchoring other cell envelope components[25–27]. The disruption of bacterial cell wall leads to cell lysis and hence cell death. Therefore, the cell wall is one of the best known and most validated targets for antibacterial therapy such as penicillin, teicoplanin, and vancomycin.

Vancomycin inhibits the transpeptidase reaction that occurs as one of the final stages of PG biosynthesis by binding to terminal D-alanyl-D-alanine residues of NAG/NAM peptides, preventing their cross-linking [19, 28]. In figure 4A and 4B, we indicated that cell wall changes such as cell wall organization, peptidoglycan-based cell wall biogenesis, and peptidoglycan metabolic and catabolic processes, were enriched significantly in Q1, but were rarely enriched in Q4. According to this result, there was a great impact on the cell wall of low level resistant *S.aureus* 11Y compared to the isogenic strain 11. Therefore, drugs interfering with the PG biosynthesis could be effective antibacterial agents. The basic architecture of PG is similar in majority of bacterial world for any molecule to act as a drug target, it is imperative that inactivation or absence of the target should result in cell death.

Besides these changes in cell wall, our data also indicated that some proteins rela ted to sugar metabolism changed obviously inside the cell. In figure 3A, the enrichment fold of glycosaminoglycans (GAGs) was 4.13. In the biological process of Figure 4A, GAG metabolic and catabolic processes were enriched in Q1. GAGs, previously called mucopolysaccharides, are linear polysaccharides composed of repeating disaccharid e units that consist of a uronic acid or galactose and a hexosamine [29]. The extraordinary structural diversity of GAGs enab-lescells to interact with a wide variety of patho gens, for instance viruses, bacteria, parasites,and fungi. GAGs-pathogen interactions ar e implicated in many steps of pathogenesis, includin-g cell adhesion and invasion, cell proliferation and differentiation, blood coagulation, and infl-ammation. It is reported tha t both Gram- negative and Gram- positive bacteria contain polysaccharid-e capsules that mi mic the structure of GAGsMicrobial GAGs are apparently used for one or more bac-terial def ence strategies including molecular mimicry, hijacking biological pathways, or altering hos-t d efences. These capsules can serve as molecular camouflage for commensal populations o f pathoge-nic bacteria allowing these microorganisms to evade the host immune response[30–32]. The results showed that GAGs may become a larger and more validated class of therapeutic targets in bacterial infection.

In addition, aspartate family amino acid metabolic and biosynthetic processes were also enriched in Q1 in Figure 4A. Recently, it was reported that different concentrations of aspartate (Asp) inhibited S. aureus biofilm formation on tissue culture plates and attenuated the cellular metabolic activity of S. aureus [33]. In our previous study, with the rise of vancomycin resistance level, S. aureus was always accompanied by a decline in the ability of biofilm formation (data not shown). Therefore we speculated that Asp plays a role in the relationship between vancomycin resistance and biofilm formation in *S. aureus*.

Via adopting the advanced TMT labeling and HPLC fractionation technique, we also successfully identified 149 differentially expressed proteins (21 up-regulated, 128 down-regulated) in Table 1. The numbers of differentially expressed protein are much more than other researchers identified in the past studies [34]. These 149 proteins take an indispensable part in the regulatory networks of *S. aureus* with vancomycin treatment, which involve in a wide ranges of biological processes, such as cell wall metabolism, cell adhesion, pressure response, cell division. And we choose some of them to describe specifically.

### Cell wall metabolism

For any molecule to act as a drug target, it is imperative that inactivation or absence of the target should result in cell death. MurA (UDP-N-acetylglucosamine 1-carboxyvinyltransferase), in the first committed step of peptidoglycan biosynthesis, catalyzes the transfer of enolpyruvate from phosphoenolpyruvate to UDP-NAG, releasing inorganic phosphate[35]. The deletion/inactivation of *murA* gene from *S. aureus* and *Streptococcus pneumoniae* has been reported to be lethal for bacteria due to the loss of cell integrity and susceptibility to osmotic lysis[36]. The importance of MurA as a drug target is also substantiated by the fact that the antibacterial activity of an antibiotic, fosfomycin is due to its covalent binding to the active site of MurA enzyme [37, 38]. MurA is neither required nor present in mammals including humans. In addition, its poor homology with mammalian proteins suggests that MurA can be a potential drug target.

### Cell adhesion

*S. aureus* serine-aspartate repeat protein C and D (SdrC, SdrD), together with fibrinogen-binding clumping factor A and B (ClfA and ClfB), are members of a structurally rela ted family of cell wall anchored proteins[39]. Both of SdrC and SdrD belong to the Sdr proteins. The Sdr proteins are members of a family of surface proteins, which are ch aracterized by the presence of an R region composed by a large stretch of repeated SD dipeptides[40] The two protein covalently anchoring to the peptidoglycan of the bac terial cell wall, are important adhesion factors of the bacteria. They regulate the bindin g capacity of *S. aureus* to fibrinogen on the host cell surface and promote bacterial in vasion into host tissues. Corrigan *et al.* reported *SdrC* and *SdrD* as well as *ClfB* each contributed to the ability of *S. aureus* to adhere to human desquamated nasal epitheli al cells. Mutant lacking either of the three genes was completely defective in adherenc e [41]. Using a silkworm model, Shinya Miyazaki *et al.* demonstrated that SdrC and Cl fB contribute to the virulence of *S. aureus* in silkworms. Especially the *sdrC*-disrupted mutant had severely attenuated virulence in silkworms, indicating that SdrC plays a pro minent role in infection by S. aureus in silkworms[42]. The self-association of the serin easpartate repeat protein SdrC contribute to both bacterial adherence to surfaces and biofilm formation[43].SdrD is essential for abscess formation [44]. Trad et al. indicat ed that the widespread existence of sdrD gene in S.aureus was responsible for bone i nfections[45]. Sabat et al. also reported that the sdrD gene was significantly associated with osteomyelitis [46]. Josefsson et al. elucidated that ClfA is indispensable during th e early stage of infection, and its virulence characteristic was conformed by experiment al animal infection models[47–50]. In general, the four surface proteins are involved in t he adhesion of *S. aureus* and theirdifferent expression levels will have a significant im pact on the ability of *S.aureus* pathogeni-city.In addition, another study has shown tha t, in *S. aureus,* Eep2 can promotes bacterial adhe-sion to a human epithelial cell line through regulating the expression of adhesins.[51]. Surfac-e proteins iron-regulated surface determinant (IsdA) has been reported to be able to promote adhesi-n in vitro.The up-regulation of genes encoding surface proteinsIsdA which promote the adhesion of S. aureu s in ampicillin induced biofilm explained the enhanced biofilm viability and biomass[52].

### Proteolysis

Atl is the most predominant peptidoglycan hydrolase in staphylococci. It is a bifunctional enzyme that possess both amidase and glucosaminidase activities[53]. Atl has various functions. During cell division, this autolysin is responsible for hydrolyzing the amide bood between N-acetylmuramic acid and l-alanine[54]; it acts directly as an adhesin by binding to fibronectin, fibrinogen and vitronectin [55]; it is involved in impairment biofilm formation [56]; and finally, *atl* mutants were attenuated in pathogenesis in an intravascular catheter-associated rat infection model[54]. Atl and LytM have been shown to have enzymatic activities reducing muropeptide cross-linkage, glycan chain length and/or cell division in S. aureus.LytM has been described as a cell wall gly-cyl-glycine endopeptidase cleaving the S. aureus muropeptide cross bridge. SceD was previously identified in the extracellular fraction of S. aureus and has a lysozyme-like transglycosylase motif according to the Pfam motifdatabase (www.sanger.ac.uk/cgi-bin/Pfam). Similar abundance changes for LytM and SceD were found at the gene expression level.The transglycolase SceD is also an important autolysin in *S. aureus.* It was up-regulated in both linezolid-resistant and daptomycin-resistant strains[56]. In our study, SceD expressed highly in VISA strain 11Y compared to MSSA strain11Y, its expression ratio was 14.084. The abundance of autolysins in vancomycin resistance *S. aureus* suggests some roles for these proteins in resistance against vancomycin and probably other cell wall inhibitors.

### Pressure response

Methionine sulfoxide reductases (Msr) are important protein repair enzymes that catalyze the reduction of methionine sulfoxide (Met-O) to methionine[57]. They play a pivotal defensive role against oxidative stress from bacteria to humans. There are basically two types, referred to as MsrA and MsrB. *S. aureus* contains three genes encoding MsrA (MsrA1, MsrA2 and MsrA3) and an additional gene that encodes MsrB[57]. In our experiment we identified MsrA2 and MsrB. Both MsrA2 and MsrB have a key function as a repair enzyme for proteins inactivated by oxidation. In *Pseudomonas aeruginosa*, inactivation of either *msrA* or *msrB* or both increased its killing by oxidants [57].Both MsrA and MsrB contributed to the enzymatic defenses of *Mycobacterium tuberculosis* from reactive oxygen species [57]. Mutation in the *msrA* or *msrB* gene in *Enterococcus faecalis* resulted in increased sensitivity to H_2_O_2_. In addition, an *msrA-msrB* double mutant showed further increase in sensitivity suggesting that the effect of mutations were additive[57]. Although few studies have demonstrated the role of MsrA2 and MsrB in *S. aureus* resistance to oxidative stress, according to the studies mentioned above, we can speculate that increased expression of these two proteins appears to be a compensatory response used by low level resistant *S. aureus* to ensure that oxidative damage resistance is maintained.

AhpC, an alkyl hydroperoxide reductase subunit C, belongs to the alkyl hydroperoxide reductase system, which has activity against H_2_O_2_, organic peroxides, and peroxynitrite[58]. Work in Burkholderia thailandensis suggests that AhpC is the major scavenger of H2O2 generated in the cytoplasm of this bacterium as a

by-product of aerobic metabolism[59]. Under aerobic growth, AhpC is required to prevent cell damage in *S. aureus*. It is a major component involved in organic-hydroperoxide resistance and scavenging H_2_O_2_[60].

There are still lots of differentially expressed proteins in our Table 1, we did not describe all, but they can not be ignored in the role of bactericidal action of vancomycin. Taken all our data together, we sum up that vancomycin treatment affects *S.aureus* regulatory networks not only occurs in the external but also internal cells. It is extremely complex and involves multiple factors, such as the changes of the cell wall, amino acid mechanism, proteolysis, cell adhesion, pressure response. So we infer that resistance to vancomycin in *S. aureus* is the result of various biological processes that are mutually coordinated and interact with each other to neutralize a large amount of damage caused by antibiotics. And antibiotic resistance requires a fitness cost. In conclusion, taking the advantages of TMT labeling, HPLC Fractionation and LC-MS/MS analysis, quantitative comparison of the proteome, with a certain concentration vancomycin treatment was extensively studied. This provides us with a valuable understanding pertinent to vancomycin resistance in *S. aureus.* By using proteomics techniques to compare the protein composition of cross drug-resistant, bacteria, it is possible to discover their common features, thus providing a basis for the development of broad-spectrum and highly effective antimicrobial agents. These data may assist with the development of novel strategies for the prevention or treatment of infections due to this problematic organism.

### Discussion

In general, the findings mean that a better understanding of virulence factors and resistance mechanisms to antibiotics is achievable through realizing the functionalities of the involved proteins.the integration of strategies to follow the dynamic localization of proteins inside the cell gives us more information about the spatial reorganization of the cell proteome when an infection takes place.reveal bacterial resistance and virulence mechanisms, and significant new targets for future drug discovery. The immense potential of proteomic technologies to achieve a deeper insight into pathogenesis and develop therapeutic techniques is undeniable.

## Acknowledgment

The authors would like to thank professor Michael Tunney for his contribution to this study. This works was financially supported by Liaoning Natural Science Fund Grant Number: (110/1210200210).

